# COVIEdb: A database for potential immune epitopes of coronaviruses

**DOI:** 10.1101/2020.05.24.096164

**Authors:** Jingcheng Wu, Wenfan Chen, Jingjing Zhou, Wenyi Zhao, Shuqing Chen, Zhan Zhou

## Abstract

2019 novel coronavirus (2019-nCoV) has caused large-scale pandemic COVID-19 all over the world. It’s essential to find out which parts of the 2019-nCoV sequence are recognized by human immune system for vaccine development. And for the prevention of the potential outbreak of similar coronaviruses in the future, vaccines against immunogenic epitopes shared by different human coronaviruses are essential. Here we predict all the potential B/T-cell epitopes for SARS-CoV, MERS-CoV, 2019-nCoV and RaTG13-CoV based on the protein sequences. We found YFKYWDQTY in ORF1ab protein, VYDPLQPEL and TVYDPLQPEL in spike (S) protein might be pan-coronavirus targets for vaccine development. All the predicted results are stored in a database COVIEdb (http://biopharm.zju.edu.cn/coviedb/).

## Introduction

Two coronaviruses—severe acute respiratory syndrome coronavirus (SARS-CoV) and Middle East respiratory syndrome coronavirus (MERS-CoV) have caused two large-scale pandemics in the past two decades [1,2]. Now, the third coronavirus caused pandemic (COVID-19) is ongoing [3,4]. The 2019 novel coronavirus (2019-nCoV) which was first identified in Wuhan, China in December 2019, from patients with pneumonia is the very coronavirus [5]. Analysis of the viral genome has revealed that 2019-nCoV is phylogenetically close to SARS-CoV [6], as was named SARS-CoV-2. As of April 27, 2020, 2,980,053 people have been confirmed COVID-19, including 206,569 deaths (~6.9% fatality rate) all over the world.

Because of the less cost-effective than treatment, and reduce morbidity and mortality without long-lasting effects, vaccines are the most effective strategy for preventing infectious diseases [7]. However, there is still no approved vaccines for human coronaviruses (hCoV). There are several types of vaccines are under pre-clinical testing or clinical trials including inactivated vaccine, recombinant subunit vaccine, recombinant vector vaccine, and nucleic acid vaccine. In general, modern vaccines, such as recombinant subunit, peptide, and nucleic acid vaccines, are advantageous over other types of vaccines because of higher safety and less side effect, by inducing the immune system without introducing whole infectious viruses [8]. Nucleic acid vaccines such as DNA vaccines and mRNA vaccines represent an innovative approach by direct injection of plasmids or mRNAs encoding the antigens, accompanied with a wide range of immune responses [9,10]. These advantages are applied with prophylactic vaccines and therapeutic vaccines to treat infectious diseases and cancers. For the development of modern vaccines, it is of critical importance to identify potential immune epitopes of 2019-nCoV, as well as other infectious pathogens.

Considering the seriousness of the recent outbreaks of zoonotic coronaviruses, therapeutic agents and vaccines for pan-coronaviruses should be developed to cope with the hCoV outbreaks in the present and in the future. Here we predict all the potential B/T cell epitopes for SARS-CoV, MERS-CoV and 2019-nCoV to provide potential targets for pan-coronaviruses vaccine development. The prediction are based on their proteins sequences. RaTG13-CoV is added because of its high homology with 2019-nCoV (96% whole genome identity [11]). All the predicted results are stored in a database named COVIEdb (http://biopharm.zju.edu.cn/coviedb/).

## Database content and usage

### Data Source

The protein sequences of SARS-CoV, MERS-CoV, 2019-nCoV, and RaTG13-CoV are downloaded from NCBI (https://www.ncbi.nlm.nih.gov/). The genome sequences of SARS-CoV, MERS-CoV, 2019-nCoV and RaTG13 encode 13, 10, 11 and 10 proteins, respectively. Among these protein sequences, ORF1ab is not divided. And the N protein of SARS-CoV is divided into ORF9a and ORF9b.

The human leukocyte antigen (HLA) alleles used for T-cell epitopes prediction are derived from Zhou *et.al.* which analyzed the HLA distribution of 20,635 individuals of Han Chinese ancestry [12]. The top 20 HLA I alleles of A, B, C subtypes and HLA II alleles with frequency more than 5% are the final HLA datasets (Table S1).

### B-cell epitopes prediction

The B-cell epitopes were predicted by the seven tools embedded in the Immune Epitope Database and Analysis Resource (IEDB) [13]. More specifically, BepiPred-1.0 [14], BepiPred-2.0 [15], Chou and Fasman beta turn prediction [16], Emini surface accessibility scale [17], Karplus and Schulz flexibility scale [18], Kolaskar and Tongaonkar antigenicity scale [19] and Parker hydrophilicity prediction [20] are used for predicting amino acid sites belonging to B-cell epitopes. The parameters are all set as default. The sites confirmed by at least five tools would be labeled in red. The thresholds of each tool are in Table S2a. In this study, only amino acids that be confirmed by at least five tools are considered as B-cell epitopes.

### T-cell epitopes prediction

The T-cell epitopes prediction were divided into two parts. One of them are presented by HLA I allele and would induce the activation of CD8+ T cells. This type of T-cell epitopes were predicted by NetMHCpan 4.0 [21], MHCflurry [22] and DeepHLApan [23]. Another type of T-cell epitopes presented by HLA II alleles were predicted by MixMHC2pred [24] and NetMHCIIpan [25], which would induce the activation of CD4+ T cells. The thresholds to defined potential T-cell epitopes of each tool are listed in Table S2b.

In the prediction of T-cell epitopes presented by HLA I alleles, all peptides with length range from 8 to 11 were selected and combined with previous HLA I alleles to create HLA-peptide pairs. It’s similar to predict that presented by HLA II alleles, with the difference that peptide length range from 15 to 28. Only HLA-peptide pairs satisfied with all thresholds of used tools would be potential T-cell epitopes in this study.

### Web interface

The web interface of COVIEdb is made up of five parts, “Home”, “B-epitope”, “T-epitope”, “Peptide”, and “Help”.

A statistical table about the predicted B-cell epitopes and T-cell epitopes are embedded in the “Home” page. The table shows the coronavirus, gene, protein length, the number of amino acids that predicted to be potential B-cell epitopes and T-cell epitopes (Table 1). The number of the predicted epitopes are different but similar among four coronaviruses. The N protein of SARS-CoV is represented as ORF9a and ORF9b as previous state. We can find that many peptides have the potential to be B-cell epitopes or T-cell epitopes, but the prefer targets should be shared by different coronaviruses and we will discuss in the next section.

**Table 1.**
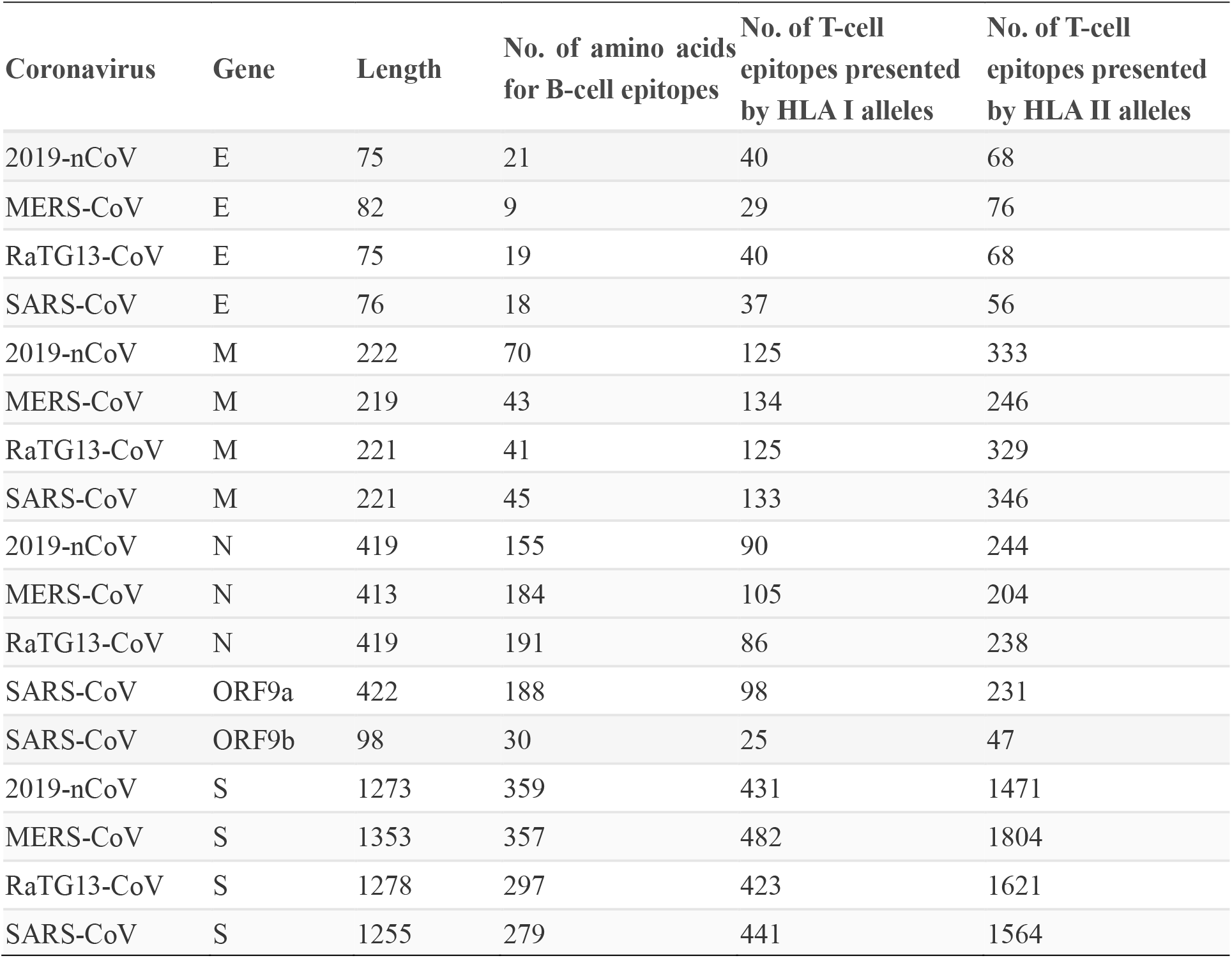
The potential immune epitopes from E, M, N, and S proteins for each coronavirus in the database.

The predicted results of B-cell epitopes could be searched in “B-epitope” page. With the virus and gene selected, the corresponding predicted B-cell epitopes would appear (Figure 1a is the example result when the selected coronavirus and protein are 2019-nCoV and S, respectively). The first to fourth columns are the basic information of the amino acids in coronavirus. The following seven columns are the results that different tools predicted if it’s the amino acid in B-cell epitopes. Once click the blue number, the corresponding score and threshold would show (Figure 1b). The last column is the number of tools that the amino acid is predicted as potential B-cell epitopes. On click the number, the corresponding tools would show (Figure 1c).

**Figure 1.**
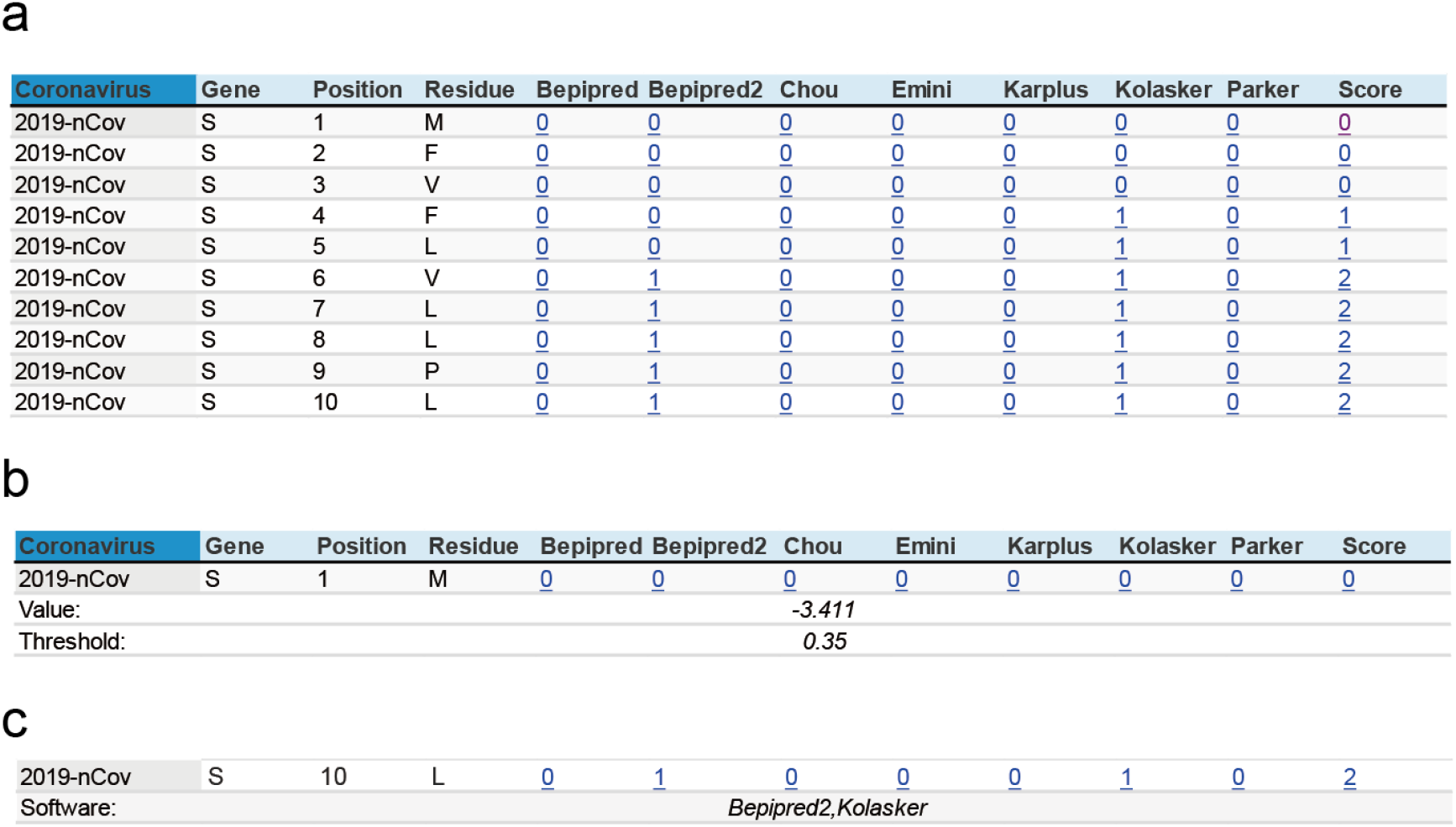
The searching result of “B-epitope” page when selecting 2019-nCoV and S. **a.** the whole interface. **b.** the predicted score and corresponding threshold on click of blue number in the column of seven tools. **c.** the detailed tools that convince the amino acids is part of B-cell epitopes.

The predicted results of T-cell epitopes could be searched in “T-epitope” page. Similar with that in “B-epitope” page, coronavirus and protein are necessary. Besides, the type of T-cell epitopes should also be selected. Because of the numerous pairs of HLA-peptide, we don’t show predicted score of all pairs. Only the peptide-HLA pairs which satisfied all thresholds would be showed in this page. Figure 2a and Figure 2b show the searching results when selecting “2019-nCoV, S and HLA I allele”, “2019-nCoV, S and HLA II allele”, respectively.

**Figure 2.**
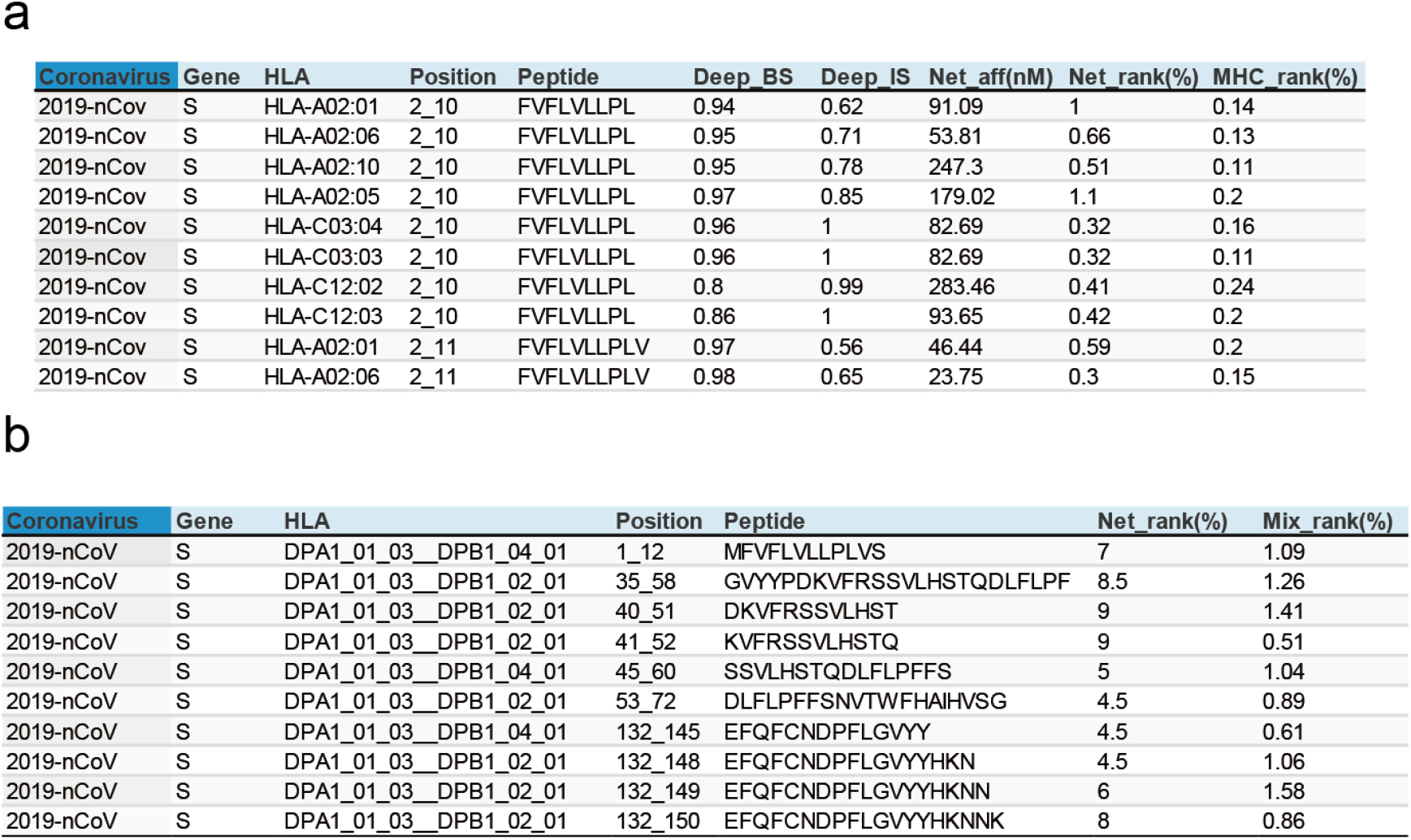
**a.** The searching result of “T-epitope” page when selecting 2019-nCoV, S and HLA-I allele. **b.** The searching result of “T-epitope” page when selecting 2019-nCoV, S and HLA-II allele.

The searchable data in the “Peptide” page is the combined result of previous predicted B-cell epitopes and T-cell epitopes. In this page, the only selectable parameter is the protein. On click the search button with protein S selected, a table will show (Figure 3a). The first column is the protein name. The second column is the peptide that either be predicted T-cell epitopes presented by HLA I alleles (with green background) or HLA II alleles (with yellow background). Amino acids that have been predicted as B cell epitopes by at least five tools are labeled in red. The third is an adjustment score to evaluate the possibility of this peptide to be B cell epitopes. The formulation is as follows:

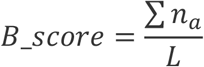

**Figure 3.**
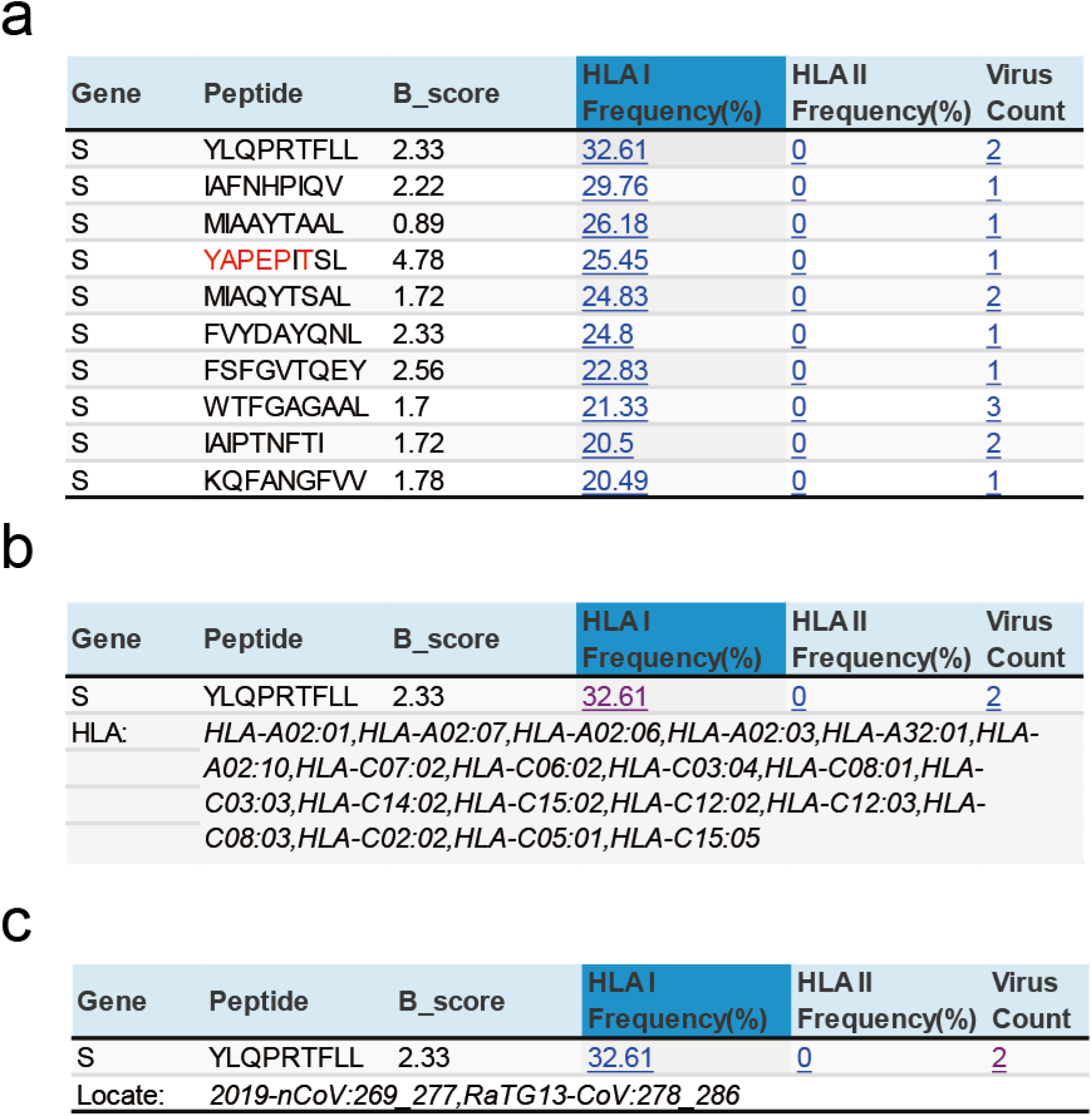
**a.** The searching result of “Peptide” page when selecting S. **b**. the detailed HLA alleles that could present peptides. **c.** The detailed coronavirus and the peptide location in this coronavirus when click the number in the last column.

Where *L* is the length of the peptide, *a* is the amino acid that belongs to the peptide, *n_a_* is the number of tools convinced that amino acid is part of B-cell epitopes. The fourth/fifth columns are the frequency of HLA I/II alleles that could present the peptide. On click the number would show the detailed HLA alleles (Figure 3b). The last column is the number of coronavirus that encodes the peptide. The detailed coronavirus and the peptide location in this coronavirus would show on click the number (Figure 3c). The table displays in descent according to the frequency of HLA I alleles by default. However, other columns could also be sortable.

### Shared B/T-cell epitopes

Though the evolution rate of human coronavirus is fast, we try to find out B/T cell epitopes conserved and shared in different coronaviruses for the pan-coronavirus vaccine development. Based on the predicted B-cell epitopes and T-cell epitopes, we found 77 peptides that exist in all coronaviruses have the potential to induce T-cell activation and 10 of them with B_score larger than 4 (Table S3, Table 2). In particular, the peptide YFKYWDQTY from ORF1ab could be presented by 7.33% people, which might be a good candidate for vaccine design.

**Table 2.** The potential T-cell epitopes with B_score larger than 4.

| Gene | Peptide | HLA able to present peptide | Peptide location in different coronaviruses | B_score | HLA I frequency | HLA II frequency |
| --- | --- | --- | --- | --- | --- | --- |
| ORF1ab | KPGGTSSGDATTAYA | DQA1_05_05__DQB1_03_01 | SARS-CoV:5045_5059 | 4.98 | 0 | 0.42 |
|  |  |  | 2019-nCoV:5068_5082 |  |  |  |
|  |  |  | MERS-CoV:5054_5068 |  |  |  |
|  |  |  | RaTG13-CoV:5067_5081 |  |  |  |
| ORF1ab | TSSGDATTAY | HLA-B15:01 | 2019-nCoV:5072_5081 | 4.98 | 1.68 | 0 |
|  |  |  | RaTG13-CoV:5071_5080 |  |  |  |
|  |  |  | SARS-CoV:5049_5058 |  |  |  |
|  |  |  | MERS-CoV:5058_5067 |  |  |  |
| ORF1ab | SSGDATTAY | HLA-B15:01 | 2019-nCoV:5073_5081 | 4.97 | 3.4 | 0 |
|  |  | HLA-B35:01 | RaTG13-CoV:5072_5080 |  |  |  |
|  |  | HLA-B15:02 | SARS-CoV:5050_5058 |  |  |  |
|  |  |  | MERS-CoV:5059_5067 |  |  |  |
| ORF1ab | KPGGTSSGDATTAYAN | DQA1_05_05__DQB1_03_01 | SARS-CoV:5045_5060 | 4.73 | 0 | 0.42 |
|  |  |  | 2019-nCoV:5068_5083 |  |  |  |
|  |  |  | MERS-CoV:5054_5069 |  |  |  |
|  |  |  | RaTG13-CoV:5067_5082 |  |  |  |
| ORF1ab | GGTSSGDATTAYAN | DQA1_05_05__DQB1_03_01 | SARS-CoV:5047_5060 | 4.7 | 0 | 0.42 |
|  |  |  | 2019-nCoV:5070_5083 |  |  |  |
|  |  |  | MERS-CoV:5056_5069 |  |  |  |
|  |  |  | RaTG13-CoV:5069_5082 |  |  |  |
| ORF1ab | GGTSSGDATTAYANS | DQA1_05_05__DQB1_03_01 | SARS-CoV:5047_5061 | 4.47 | 0 | 0.42 |
|  |  |  | 2019-nCoV:5070_5084 |  |  |  |
|  |  |  | MERS-CoV:5056_5070 |  |  |  |
|  |  |  | RaTG13-CoV:5069_5083 |  |  |  |
| ORF1ab | GTSSGDATTAYANS | DQA1_05_05__DQB1_03_01 | SARS-CoV:5048_5061 | 4.36 | 0 | 0.42 |
|  |  |  | 2019-nCoV:5071_5084 |  |  |  |
|  |  |  | MERS-CoV:5057_5070 |  |  |  |
|  |  |  | RaTG13-CoV:5070_5083 |  |  |  |
| ORF1ab | GGTSSGDATTAYANSV | DQA1_05_05__DQB1_03_01 | SARS-CoV:5047_5062 | 4.19 | 0 | 0.42 |
|  |  |  | 2019-nCoV:5070_5085 |  |  |  |
|  |  |  | MERS-CoV:5056_5071 |  |  |  |
|  |  |  | RaTG13-CoV:5069_5084 |  |  |  |
| ORF1ab | YFKYWDQTY | HLA-B15:01 | 2019-nCoV:4678_4686 | 4.08 | 7.33 | 0 |
|  |  | HLA-B15:02 | RaTG13-CoV:4677_4685 |  |  |  |
|  |  | HLA-C07:02 | SARS-CoV:4655_4663 |  |  |  |
|  |  |  | MERS-CoV:4664_4672 |  |  |  |
| ORF1ab | MGWDYPKCDR | HLA-A31:01 | 2019-nCoV:5007_5016 | 4.07 | 1.21 | 0 |
|  |  |  | RaTG13-CoV:5006_5015 |  |  |  |
|  |  |  | SARS-CoV:4984_4993 |  |  |  |
|  |  |  | MERS-CoV:4993_5002 |  |  |  |

All the T-cell epitopes shared in four coronaviruses are located in ORF1ab. However, the S protein of the coronavirus is the most important protein where the receptor binding domain (RBD) located. So we further investigated the shared epitopes that located in S protein. There are 265 potential epitopes in S protein shared by three coronaviruses and 35 of them with B_score larger than 5 (Table S4). The peptides VYDPLQPEL and TVYDPLQPEL even have B_score larger than 6, and they can be presented by 8.26% and 2.44% people, respectively, which indicates their feasibility to be pan-coronavirus vaccine targets.

The shared B/T-cell epitopes could be found in the “Peptide” page in COVIEdb and we believe that these results could benefit the vaccine development currently on 2019-nCoV and also the potential coronavirus outbreak in the future.

## Discussion

Research groups all over the world are trying to find the way to control 2019-nCoV. Several studies focus on the B/T-cell epitopes for accelerating vaccine development. The group who maintain the database IEDB identified potential targets for immune responses to the 2019-nCoV by sequence homology with closely related SARS-CoV and by a priori epitope prediction using bioinformatics approaches [26]. A recent accepted work executed a comprehensive *in silico* analysis of viral peptide-HLA I allele binding affinity across 145 HLA -A, -B, and -C genotypes for all 2019-nCoV peptides to discover the HLA susceptibility map for 2019-nCoV [27].

However, the vaccine development can’t react immediately to control a novel coronavirus epidemic. The better choice is to prepare candidate vaccines before the outbreak coming. For this purpose, we predicted all the potential B/T-cell epitopes of SARS-CoV, MERS-CoV, 2019-nCoV and RaTG13-CoV and want to find out the shared B/T-cell epitopes among them, which may be potential pan-coronavirus vaccine targets. Based on the predicted results by different tools, we find several peptides that could be vaccine candidates for hCoVs such as YFKYWDQTY in ORF1ab, VYDPLQPEL and TVYDPLQPEL in S protein. We believe that these results and the developed database could benefit not only the vaccine (especially the polyvalent vaccine which could protect from various coronavirus) development but also provide the targets for drug design such as neutralizing antibody on 2019-nCoV and the possible coronavirus outbreak in the future.

## Acknowledgment

This work was supported by the Key R&D Program of Zhejiang Province (Grant No. 2020C03010), the National Natural Science Foundation of China (Grant No. 31971371), the Zhejiang Provincial Natural Science Foundation of China (Grant No. LY19H300003), and the Fundamental Research Funds for the Central Universities of China.

